# TATA and paused promoters active in differentiated tissues have distinct expression characteristics

**DOI:** 10.1101/2020.07.15.196493

**Authors:** Vivekanandan Ramalingam, Malini Natarajan, Jeff Johnston, Julia Zeitlinger

## Abstract

Core promoter types differ in the extent to which RNA polymerase II (Pol II) pauses after initiation, but how this difference affects their tissue-specific gene expression characteristics is not well understood. While promoters with Pol II pausing elements are active at all stages of development, TATA promoters are highly active in differentiated tissues. We therefore used a genomics approach on late-stage *Drosophila* embryos to analyze the properties of promoter types. Using tissue-specific Pol II ChIP-seq, we found that paused promoters have high levels of paused Pol II throughout the embryo, even in tissues where the gene is not expressed, while TATA promoters only show Pol II occupancy when the gene is active. This difference between promoter types is associated with different chromatin accessibility in ATAC-seq data and different expression characteristics in single-cell RNA data. The results suggest that promoter types have optimized different promoter properties: paused promoters show more consistent expression when active, while TATA promoters have lower background expression when inactive. We propose that tissue-specific effector genes have evolved to use two different strategies for their differential expression across tissues.

## Introduction

The core promoter is the ~100 bp-long sequence surrounding the transcription start site (TSS) of genes that facilitates the assembly of the transcription machinery and Pol II transcription (Haberle and Stark 2018; Smale and Kadonaga 2003). Pol II transcription may be stimulated in a tissue-specific manner by activation signals from enhancer sequences (Spitz and Furlong 2012; Banerji et al. 1981), but a core promoter may also produce basal or background levels of transcription in the absence of an activation signal (Verrijzer and Tjian 1996; Kim et al. 1994; Juven-Gershon et al. 2006). Ideally, a promoter produces only minimal background expression in this inactive state, is highly responsive to enhancers and reliably produces the desired level of transcription in the active state.

A core promoter element that strongly promotes Pol II initiation is the TATA box (Patikoglou et al. 1999; Reeve 2003), an ancient core promoter element present in archaea, fungi, plants, and animals (Reeve 2003; Patikoglou et al. 1999). TATA is bound by TATA-binding protein (TBP) and helps assemble the pre-initiation complex (Nikolov et al. 1992; Kim et al. 1993; Patikoglou et al. 1999). After initiating transcription, Pol II may then pause 30-50 bp downstream of the TSS, before being released into productive elongation (Adelman and Lis 2012).

Core promoter elements not only influence Pol II initiation, but also Pol II pausing. Promoters with very stably paused Pol II are enriched for downstream pausing elements such as the Pause Button (PB) and the Downstream Promoter Element (DPE) (Hendrix et al. 2008; Lim et al. 2004; Burke and Kadonaga 1997; Shao and Zeitlinger 2017; Gaertner et al. 2012). Swapping core promoter elements or the entire promoter alters the amount and duration of Pol II pausing in *Drosophila* (Lagha et al. 2013; Shao et al. 2019).

The amount of Pol II pausing at a promoter appears to influence the expression characteristics. Promoters with high occupancy of paused Pol II are prevalent among genes that are highly regulated during development (Zeitlinger et al. 2007; Muse et al. 2007; Gaertner et al. 2012) and mediate more synchronous gene expression between cells (Boettiger and Levine 2009; Lagha et al. 2013). Without well-timed gene activation, coordinated cellular behaviors such as gastrulation may not proceed properly (Lagha et al. 2013).

The expression characteristics of TATA promoters in developing embryos are less understood. Across metazoans, genes with TATA elements are particularly enriched among effector genes, the genes responsible for the structure and function of differentiated tissues (Schug et al. 2005; Lenhard et al. 2012; Carninci et al. 2006; Engström et al. 2007; FitzGerald et al. 2006; FANTOM Consortium and the RIKEN PMI and CLST (DGT) et al. 2014). Effector genes start to be expressed when cells begin differentiation into morphologically distinct tissues (Erwin and Davidson 2009), which occurs at later stages of embryogenesis. These stages are typically not well studied, and thus the reason for the prevalence of TATA promoters in the expression of effector genes is not clear.

TATA promoters may confer different expression characteristics on effector genes. TATA promoters are often associated with higher expression variability (Tirosh et al. 2006; Lehner 2010; Hornung et al. 2012; Blake et al. 2006; Raser and O’Shea 2004; Sigalova et al. 2020). Furthermore, TATA promoters, which are expressed in the very early embryo before cellularization and pattern formation, show minimal Pol II pausing (Chen et al. 2013). Later expressed TATA genes also appear to have altered pausing behaviors, but here the evidence is conflicting and is predominantly based on cultured cells from late embryos (Shao and Zeitlinger 2017; Gilchrist et al. 2010; Krebs et al. 2017).

Here, we systematically analyzed the relationship between promoter types and gene expression in differentiated tissues of the late *Drosophila* embryo, where both TATA and paused promoters are active. We mapped the gene expression programs of all cell types using single-cell RNA-seq (scRNA-seq) and determined the occupancy of Pol II in a tissue-specific fashion. Our analysis revealed large differences in Pol II pausing between the promoters of different effector genes and showed that TATA promoters are strongly enriched among effector genes with minimal Pol II pausing. Notably, scRNA-seq revealed that TATA genes have higher expression variability but lower background expression than paused promoters and that this property correlates with lower chromatin accessibility. We propose that different promoter types are optimal for different expression properties and discuss the mechanisms by which these differences in promoter function occur.

## Results

### Characterization of the tissue-specific expression programs in the late Drosophila embryo using single-cell RNA-seq

To obtain an unbiased global view of the gene expression programs in differentiated tissues, we performed scRNA-seq on dissociated cells from late *Drosophila* embryos (Figure 1A). We chose embryos at stage 16 (14h-14.5h after egg deposition) when the tissues are fully formed but the outside cuticle is not yet developed enough to hamper the dissociation of the cells. After processing the cells through a 10× Genomics Chromium instrument (Zheng et al. 2017; Klein et al. 2015; Macosko et al. 2015), we obtained the expression profiles of approximately 3,500 cells. Cells prepared and sequenced from two separate batches yielded results that were highly similar with regard to data quality and results from clustering (Supplementary Fig. S1).

**Figure 1:**
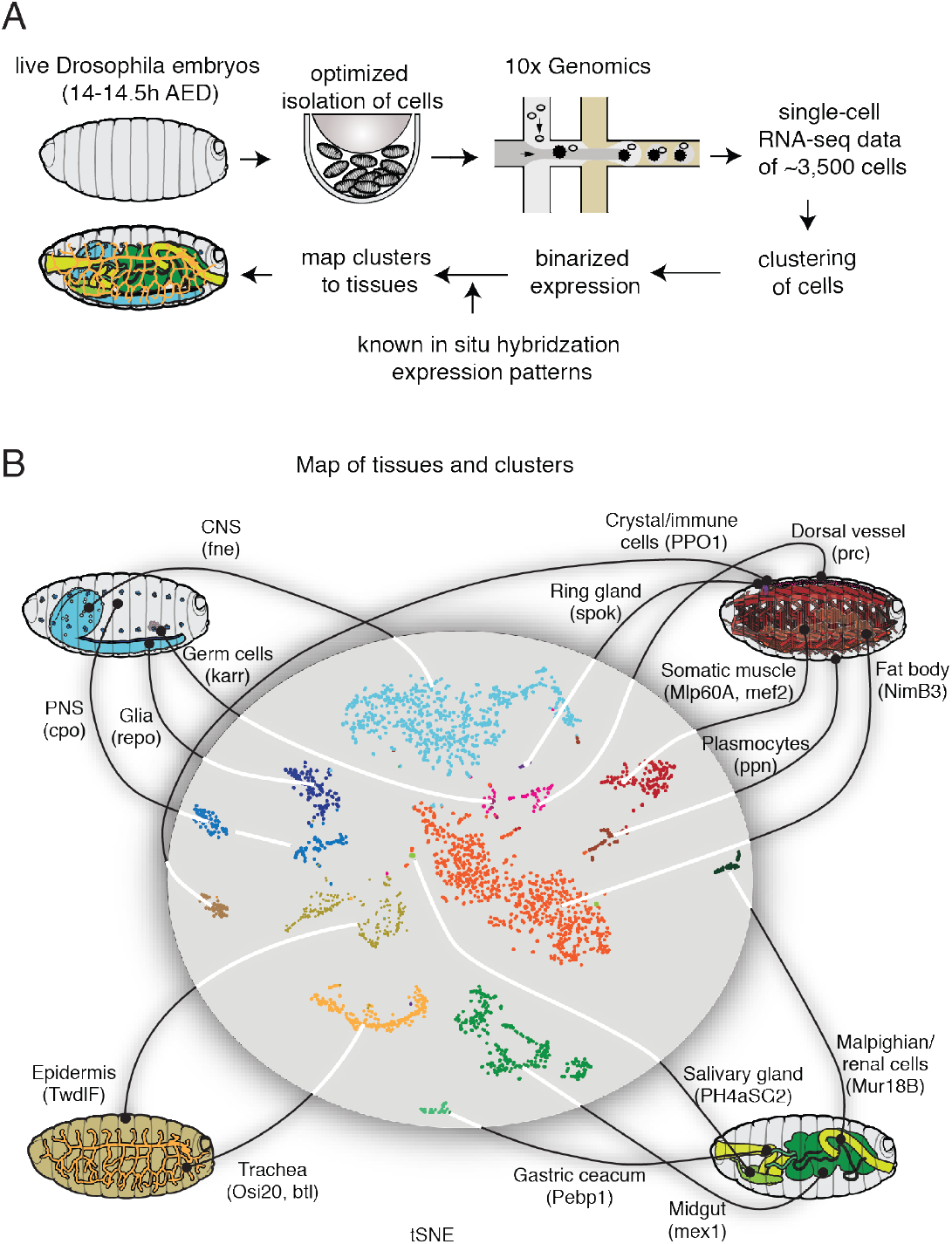
scRNA-seq captures the expression profiles of effectors genes in the late stages of *Drosophila* embryogenesis. (A) Single cells were isolated from *Drosophila* embryos 14-14.5h after egg deposition (AED). Isolated cells were processed through a 10× Genomics instrument. After sequencing the resulting libraries, the reads were aligned and processed using the standard pipeline from 10× Genomics. The single-cell gene expression profiles were used to map the cells to known cell types by comparing against the available *in situ* hybridization patterns from the Berkeley *Drosophila* Genome Project. (B) A tSNE projection of the scRNA-seq data is shown in the middle, and the known tissues to which the clusters were assigned to are graphically illustrated outside. Marker genes for each tissue type are shown in parentheses.

To identify the tissues to which each scRNA-seq cluster belongs, we correlated the scRNA-seq data with the large-scale *in situ* hybridization data from the Berkeley *Drosophila* Genome Project (BDGP) (Hammonds et al. 2013; Tomancak et al. 2007; Tomancak et al. 2002) (Figure 1A). For ambiguous clusters, we analyzed the occurrence of known tissue markers and manually merged or separated clusters such that they better matched anatomical structures. In this manner, we obtained scRNA-seq data for 16 tissues of the late *Drosophila* embryo: central nervous system (CNS), peripheral nervous system (PNS), glia, germ cells, epidermis, trachea, muscle, dorsal vessel, fat body, plasmocytes, crystal cells, ring gland, salivary gland, gastric caecum, midgut and malpighian tubules (Figure 1B, Suppl. Excel 1). For each tissue, we then identified marker genes that reliably represented the cluster, some of which were previously known (Figure 1B and S2A, S2B).

### The Pol II occupancy pattern across tissues reveals two promoter types

To characterize the promoter types and expression characteristics of effector genes, we then performed Pol II ChIP-seq experiments on a variety of tissue types from the late *Drosophila* embryo. We isolated tissue-specific cells using the INTACT method, in which nuclei from a tissue of interest are genetically tagged for biotin labeling and isolated with the help of streptavidin-coupled magnetic beads (Bonn et al. 2012; Deal and Henikoff 2011) (Figure 2A). We used fixed embryos (14-17h) as starting material and isolated the following six tissues to perform Pol II ChIP-seq experiments: neurons (using elav-Gal4), glia (using *repo*-Gal4), muscle (using *mef2*-Gal4), trachea (using *btl*-gal4), and epidermis and gut (using enhancer trap-gal4 lines 7021 and 110394, respectively, see Methods) (Figure 2B).

**Figure 2:**
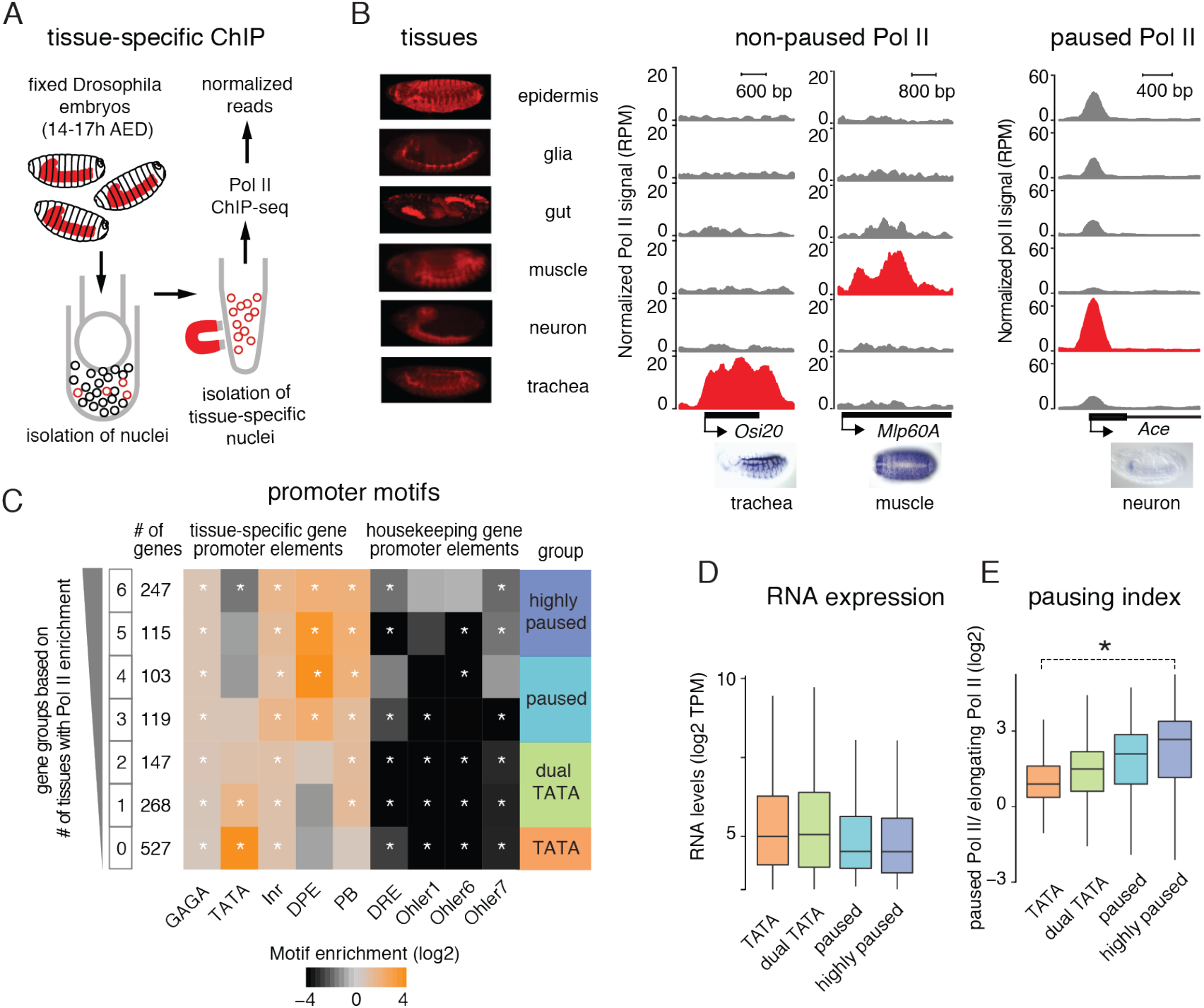
Tissue-specific Pol II ChIP-seq shows differences in Pol II occupancy patterns at effector genes. (A) Tissue-specific ChIP-seq was done by isolating nuclei from specific tissues (shown in red) by expressing the *Escherichia coli* biotin ligase (BirA) and the biotin ligase recognition peptide (BLRP) fused with a nuclear envelope-targeting sequence in the tissue of interest. This allows the isolation of nuclei from the tissue of interest using streptavidin magnetic beads. (B) Pol II ChIP-seq was performed in six different tissues shown in the left panel. The middle and the right panels show the read count normalized Pol II ChIP-seq tracks (RPM) from the six tissues at individual genes. For each gene, grey and red tracks indicate non-expressing tissues and expressing tissue, respectively. The middle panel shows the Pol II profile at two non-paused genes, which have Pol II only in the expressing tissues. The right panel shows the Pol II profile at a paused gene, which has Pol II in all observed tissues. The differences in paused Pol II occupancy at paused promoters are related to sample preparation and transcription levels. (C) Identified effector genes were grouped into seven groups based on the number of tissues in which they have Pol II enrichment above background. Pol II enrichment was calculated in a window starting from the Transcription Start Site (TSS) and ending 200bp downstream of the TSS. Distinct core promoter elements are differentially enriched in different groups (Fisher's exact test with multiple-testing correction, *P < 0.05). Only the groups with Pol II enrichment in 0 or 1 tissues are enriched for the TATA motif. Groups with TATA and pausing elements are referred to as dual. (D) RNA levels (log2 TPM) from 14-17h embryos at the different effector gene groups are shown. The TATA genes are expressed at levels comparable to the paused genes. (E) Pausing indices (log2), the ratio of Pol II at the promoter vs the gene body, are shown for the different effector gene groups. The pausing indices of genes from the TATA group are significantly lower than that of the highly paused group (Wilcoxon two-sided test, *P <10^−15^).

The Pol II ChIP-seq tracks from the six tissues confirmed that the ChIP-seq data are tissue-specific. For example, the tracheal gene *Osi20* and the muscle gene Mlp60A showed high Pol II occupancy in the trachea and muscle samples respectively, but not in the other tissues (Figure 2B middle panel). Furthermore, we compared the Pol II occupancy data with the expression programs obtained from the single-cell RNA-seq data and found an overall correlation between Pol II occupancy and gene expression (Figure S4A). This led us to conclude that the Pol II profiles are indeed from the expected tissue, and that cross-contamination was not observed.

The Pol II occupancy was however not always tissue-specific. Some genes showed high Pol II occupancy at the promoter in many or all tissues, despite being expressed in a very tissue-restricted fashion. For example, expression of *Ace* is restricted to neuronal populations, but showed very high Pol II promoter occupancy in all tissues (Figure 2B right panel). Moreover, the Pol II pattern along the gene was indicative of Pol II pausing since the Pol II occupancy peaks at the pausing position (30-50 bp downstream of the TSS) and is not detectable at substantial levels at the gene body (Figure 2B right panel). This profile of Pol II pausing is in contrast to the *Osi20* and *Mlp60A* genes, for which Pol II occupancy was tissue-specific and found throughout the body of the gene (Figure 2B middle panel).

These results suggest that there are two types of tissue-specific promoters that are regulated in a fundamentally different fashion. At one type of promoter, Pol II is recruited only in tissues where the gene is expressed and proceeds towards productive elongation without detectable pausing. On the other end of the spectrum is a promoter type where Pol II is widely recruited and found paused across all tissues, and Pol II only proceeds towards productive elongation in the tissues where the gene is expressed.

### Pol II penetrance across tissues separates TATA and paused genes

Since we had set out to analyze TATA and paused promoters, we asked what promoter types are enriched among effector genes. By eliminating developmental genes and ubiquitously-expressed housekeeping genes from all late-expressed genes, we identified a group of putative effector genes, which were enriched for GO terms of tissue-specific biological functions, e.g., synaptic transmission and chitin-based cuticle development (Suppl. Figure S3A). This group of genes was indeed enriched for TATA elements, while sequence motifs found in housekeeping genes such as the Dref Response Element (DRE) were under-represented (Suppl. Figure S3C). Besides, sequence motifs typically found in paused promoters such as DPE and PB were also significantly enriched, suggesting that effector genes are induced by both TATA and paused promoter types (Suppl. Figure S3C)

We then analyzed whether the core promoter types differ in their Pol II occupancy pattern across tissues. We grouped genes by the number of tissues (from 0 to 6 tissues) in which Pol II was detected around the transcription start site above background levels. We found that TATA genes and highly paused genes were at the two extreme ends of the Pol II penetrance pattern (Figure 2C). Promoters, where Pol II was found across most tissues, were depleted for TATA elements and were highly enriched for pausing elements such as the DPE and PB, consistent with being highly paused. Promoters with Pol II occupancy in very few tissues, on the other hand, were highly enriched for TATA (Figure 2C). Strikingly, the most canonical TATA promoters, which contain TATA in the absence of pausing elements, were found in the group with no apparent Pol II occupancy (Figure 2C), although RNA-seq data suggest that these genes are transcribed (Figure 2D). This suggests that Pol II may be hard to detect at some canonical TATA genes, presumably because Pol II does not pause, and thus significant levels can only be detected with high levels of transcription.

We conclude that the penetrance of Pol II occupancy across tissues is a measurement for Pol II pausing. Consistent with this, we found the pausing index, the ratio of Pol II at the promoter vs the gene body, to correlate with the Pol II penetrance measurements, with TATA genes showing the least amount of Pol II pausing (Figure 2E). To rule out that Pol II is observed in fewer tissues simply because the expression levels are lower and do not pass the threshold of Pol II detection, we analyzed the expression levels in these gene groups. There was no positive correlation between Pol II penetrance and transcript levels across the embryo (Figure 2D). The expression of TATA genes is initially slightly lower in embryos at 14-14.5h, but their expression rises over time and is overall higher than that of highly paused genes (Figure S5A and S5B). Thus, genes with Pol II occupancy across many tissues were overall not expressed at higher levels than those with Pol II in only a few tissues, consistent with paused Pol II being more likely associated with lower expression levels (Shao and Zeitlinger 2017).

These results suggest that effector genes are induced by different promoter types. Since effector genes have previously been associated with TATA promoters (Schug et al. 2005; Lenhard et al. 2012; Carninci et al. 2006; Engström et al. 2007; FitzGerald et al. 2006; FANTOM Consortium and the RIKEN PMI and CLST (DGT) et al. 2014), we investigated whether the putative effector genes with highly paused promoters could be functionally distinguished from the effector genes with TATA promoters. GO analysis did not reveal any clear functional differences between these groups (Figure S3B). We did notice that the TATA effector genes were often short genes found in clusters of gene families (e.g. the *Osi* gene family, Figure S6), and many of them were expressed in tissues such as the epidermis, gut, and trachea, which are exposed to the environment and may require adaptation (Dorer et al. 2003; Shah et al. 2012; Cornman 2009).

### TATA genes are expressed with high variability but low background expression

Using the scRNA-seq data, we next analyzed whether the different promoter types might display different expression characteristics across tissues. We first analyzed their expression noise, measured by the coefficient of variation across different expression bins. We found that TATA genes had consistently higher expression variation than paused genes in all expression bins (Figure 3A).

**Figure 3:**
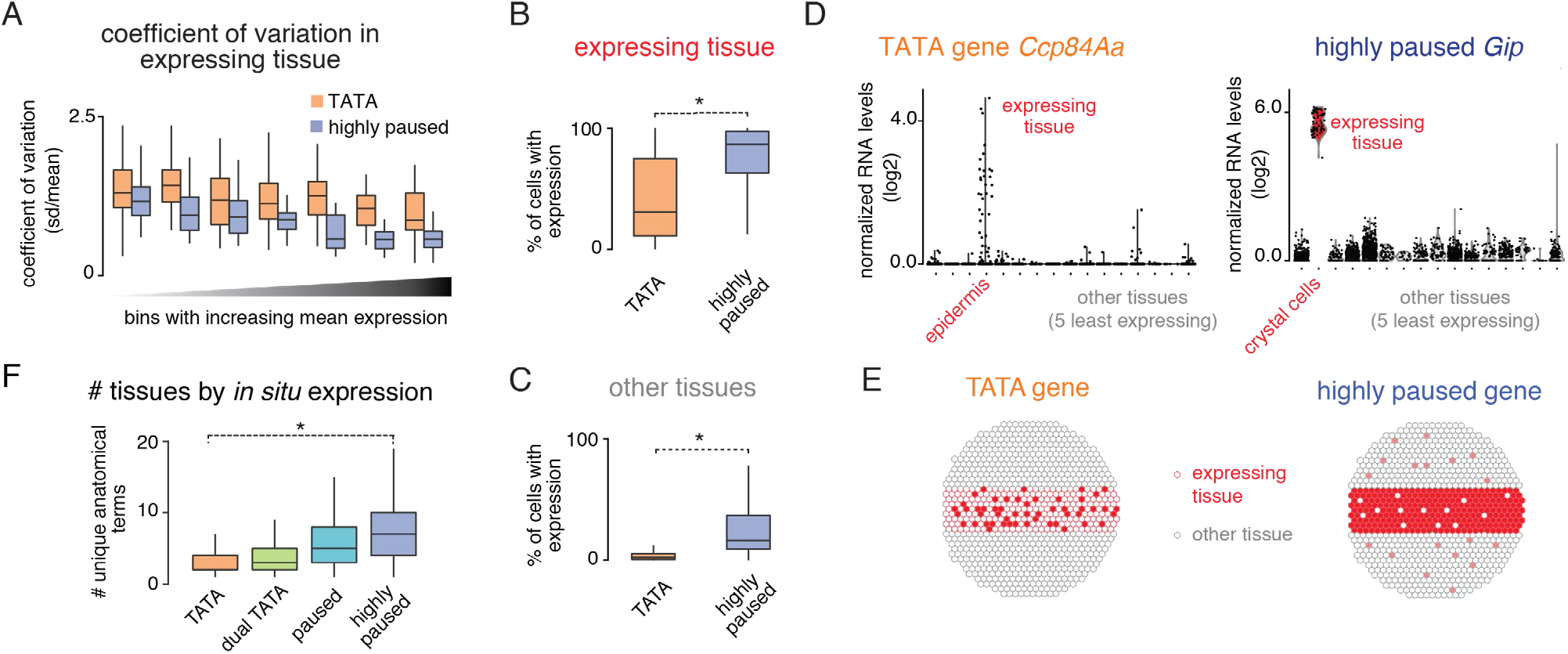
scRNA-seq reveals differences in expression characteristics of TATA and paused genes. (A) The differences in cell-to-cell gene expression variability for the TATA and the paused effector gene groups were analyzed using the scRNA-seq data. The coefficient of variation (standard deviation/mean) of gene expression was calculated for all genes in the tissue with the highest expression for each gene. The median coefficient of variation was consistently lower for the paused genes compared to the TATA genes. (B & C) The frequency of cells with any detectable expression was calculated in tissues with the highest expression for each gene (expressing tissue) and in five other tissues with the least expression for each gene (other tissues). (B) The frequency of cells with detectable expression in the expressing tissues, a measure of expression robustness, is lower for the TATA genes compared to the highly paused genes. (Wilcoxon two-sided test, *P <10^−15^). (C) The frequency of cells with detectable expression in the other tissues, a measure of background expression, is also lower for the TATA genes compared to the paused genes (Wilcoxon two-sided test, *P <10^−15^). (D) Normalized gene expression levels (read count normalized for each cell, log2) in different tissues, from the scRNA-seq experiment, for a TATA group gene, and a highly paused group gene are shown. The TATA gene, *Ccp84Aa*, shows noisy expression in the epidermis, without detectable background expression in non-expressing tissues. The highly paused gene, *Gip*, shows very robust expression in crystal cells, but has high background expression in the non-expressing tissues. (E) A schematic representation of the expression characteristics of TATA and highly paused genes is shown. Grey hexagons represent the non-expressing tissue, while red hexagons represent the expressing tissue. (F) The number of annotation terms associated with each gene in the BDGP *in situ* database. This is a measure of whether the expression of a gene is restricted to specific subsets of tissue (*P <10^−15^).

We then specifically compared the expression variability between TATA and highly paused promoters in tissues where the genes are either active or inactive. We did so by scoring how many cells had a detectable expression for each gene in the corresponding expression clusters. The results show that paused genes are indeed more consistently expressed between cells when active in a tissue, but they also have higher background expression when not expressed (Figure 3B,C). In contrast, TATA genes show higher expression variability when expressed, but less background expression in tissues where the gene is not expressed. This difference between promoter types was still present after accounting for differences in their expression levels (Figure S7A,B). This pattern is illustrated by *Ccp84Aa,* a TATA gene expressed in the epidermis, and *Gip,* a highly paused gene expressed in crystal cells (Figure 3D). A representation of the distinct expression properties of the two promoter types is shown in Figure 3E.

The high expression variability of TATA genes has previously been associated with high stochastic noise that is intrinsic to this core promoter type (Hornung et al. 2012; Blake et al. 2006; Raser and O’Shea 2004). While our results are consistent with these findings, it is also possible that TATA genes appear to be more variably expressed across cells within a tissue because they are expressed in a more restricted fashion, e.g. in subtypes of cells within a tissue. To test this, we analyzed their embryonic expression pattern in late-stage *Drosophila* embryos using the BDGP *in situ* database. We found that the TATA genes had indeed significantly fewer annotated expression patterns compared to highly paused genes (Figure 3F). This more restricted expression of TATA genes may therefore also contribute to the observed expression variability in our scRNA-seq data.

### TATA promoters have lower chromatin accessibility

We have previously observed that paused and TATA promoters in the *Drosophila* embryo have different nucleosome configurations. Highly paused promoters have a strong disposition for a promoter nucleosome that is depleted when paused Pol II is present (Gaertner et al. 2012; Chen et al. 2013; Gilchrist et al. 2010). Consistent with observations in yeast, this suggests a model in which the promoter nucleosome at TATA promoters represents a barrier to transcriptional activation, which in turn may influence its expression characteristics (Tirosh and Barkai 2008; Hornung et al. 2012; Raser and O’Shea 2004).

We therefore tested whether the promoter groups among the effector genes differed in their promoter nucleosome occupancy. Using MNase-seq data, we found that the higher the Pol II penetrance across tissues, the more the promoters became nucleosome-depleted in the late embryo as compared to the early embryo. TATA promoters did not show such a change (Figure 4A arrow). This supports the idea that the expression characteristics of the two promoter types involve different nucleosome configurations.

**Figure 4:**
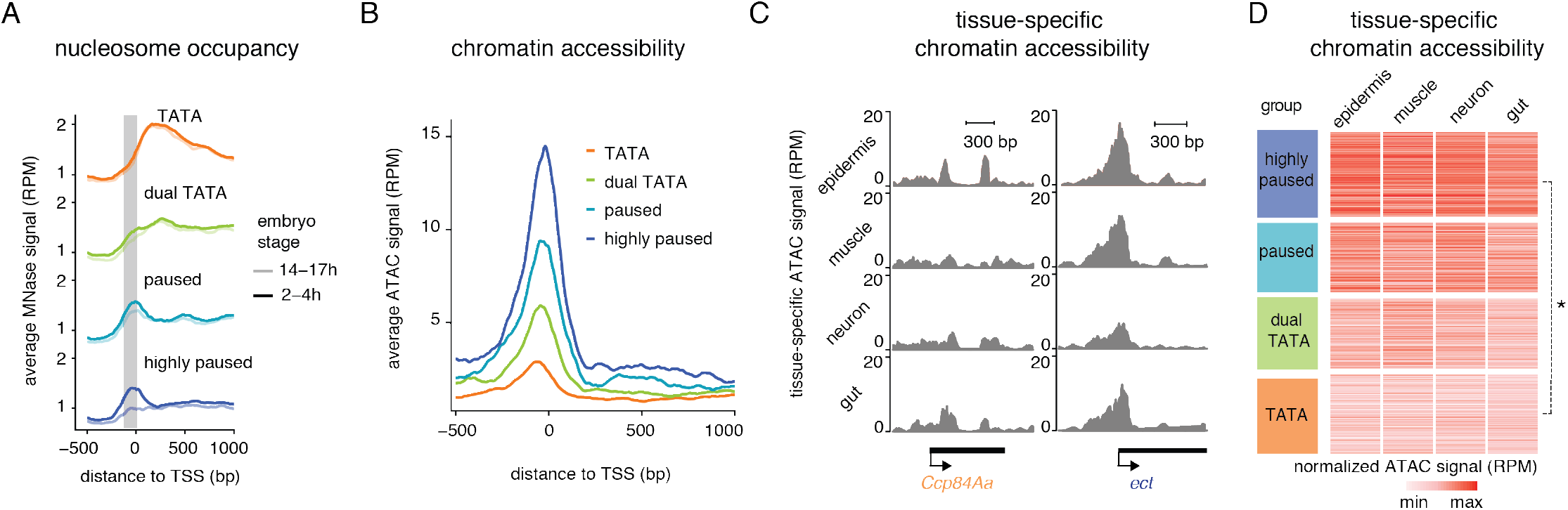
Paused genes are more accessible than the TATA genes. (A) Average read-count normalized MNase signal (RPM) from 2-4h and 14-17h embryos are shown at the different effector gene promoter groups. The grey area highlights the changes in nucleosome occupancy, which are prominent at highly paused genes (arrow). Chromatin accessibility is shown as the average read-count normalized ATAC-seq signal (RPM) from 14-17h for each promoter group. Paused genes show higher accessibility at the promoter region than the TATA genes. (C & D) ATAC-seq chromatin accessibility was measured in different tissues isolated from 14-17 h embryos using INTACT. (C) Read-count normalized ATAC (RPM) signal at individual genes from the TATA group (*Ccp84Aa*), and highly paused gene group (*ect*) are shown. (D) Read-count normalized ATAC signals (RPM) from different tissues were calculated for each gene from 150bp upstream of the TSS to the TSS. Paused genes show higher average accessibility across all tissues compared to the TATA genes (Wilcoxon two-sided test, *P <2.2*10^−16^).

If the promoter nucleosome indeed represents a barrier to activation and this barrier is lowered by the presence of paused Pol II, one would expect that promoters with paused Pol II show higher chromatin accessibility. To test this, we performed ATAC-seq experiments in 14-17h embryos as a measurement for chromatin accessibility across all tissues (Figure 4B-D). Strikingly, we found that the more Pol II pausing, the higher the average promoter accessibility, while the most canonical TATA promoters with the least amount of pausing showed the least chromatin accessibility (Figure 4B).

We then tested whether the pattern of chromatin accessibility across promoters was tissue-specific and performed ATAC-seq on tissues isolated using the INTACT method. We found that the promoter accessibility varied little between tissues (Figure 4C,D), consistent with the model that it reflects the basal state of the promoter and is not predominantly determined by the activation levels of the promoter. This supports our hypothesis that promoters with high levels of Pol II pausing are nucleosome depleted. This in turn could lower the barrier for activation, which explains the robust tissue-specific expression, but also higher levels of background expression as compared to TATA promoters (Figure 5).

**Figure 5:**
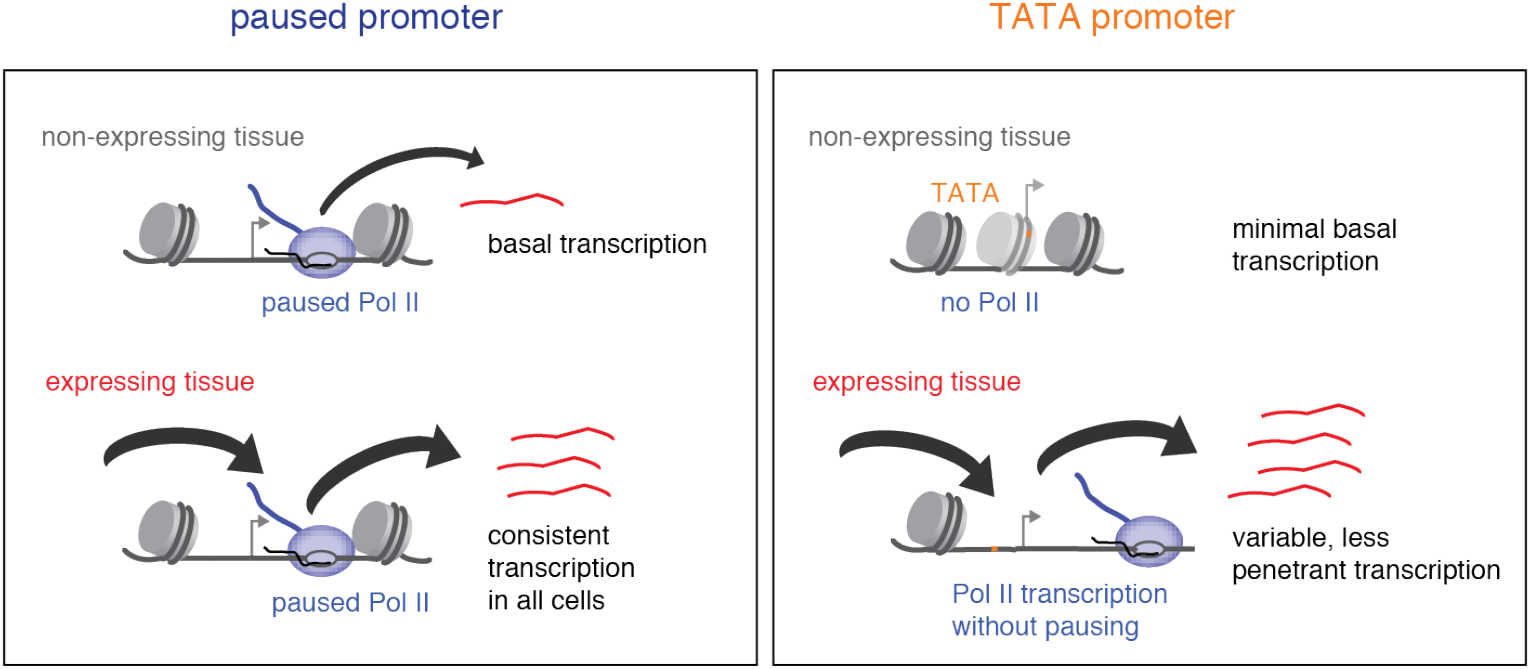
Model for the different expression characteristics of TATA versus paused promoters. Paused promoters have high levels of paused Pol II throughout the embryo, even in tissues where the genes are not expressed; they show high promoter accessibility in all tissues, low gene expression noise when active, but also high background expression when inactive. TATA promoters mediate highly tissue-restricted expression and only show Pol II occupancy when active; they have lower chromatin accessibility and background expression but show higher expression noise when active.

## Discussion

Previous bioinformatics analyses suggested that effector genes are highly enriched among genes with TATA elements (Schug et al. 2005; Carninci et al. 2006; Engström et al. 2007; FitzGerald et al. 2006; FANTOM Consortium and the RIKEN PMI and CLST (DGT) et al. 2014), but how the promoter type affects their tissue-specific expression has not been clear. Moreover, TATA genes have been observed in other contexts to have altered Pol II pausing behavior (Shao and Zeitlinger 2017; Gilchrist et al. 2010; Chen et al. 2013), but whether this applies to effector genes in differentiated tissues was not known. Furthermore, it was unclear whether effector genes would be predominantly expressed by TATA promoters or whether other promoter types would also be common for these types of genes. Here we found that effector genes with TATA promoters are indeed expressed with the least amount of Pol II pausing, but that many effector genes are also expressed from paused promoters. Since the two promoter types are employed in the same cells, we were able to directly compare the two promoter types side-by-side across different tissues, and analyze their tissue-specific expression characteristics using scRNA-seq.

We found that paused and TATA promoters regulate the tissue-specific gene expression of effector genes in fundamentally different ways. At promoters with pausing elements, paused Pol II is found broadly at promoters across tissues, even in tissues where the genes are not expressed. In contrast, TATA promoters only recruit Pol II in the tissues where the genes are active and the most canonical TATA promoters do not show Pol II pausing. These patterns of Pol II recruitment across tissues are consistent with those observed over developmental time, since we have previously observed that paused promoters typically show Pol II occupancy prior to induction, while TATA promoters do not (Gaertner et al. 2012). Our results therefore consolidate the fundamental difference between these two promoter types.

The difference in Pol II recruitment between promoter types could explain the difference in expression characteristics that we observed in our scRNA-seq data. Paused genes showed significantly lower expression variability within the cells of a tissue as compared to TATA genes. This is consistent with the more synchronous expression of paused genes over time and the generally higher expression variability of TATA genes (Tirosh and Barkai 2008; Tirosh et al. 2006; Lehner 2010; Faure et al. 2017; Sigalova et al. 2020; Hornung et al. 2012; Blake et al. 2006; Raser and O’Shea 2004; Day et al. 2016; Boettiger and Levine 2009; Lagha et al. 2013). However, paused genes also had higher levels of background expression than TATA genes.

We therefore propose that the presence of paused Pol II, while allowing low variability in gene expression, causes background expression, because of the broad recruitment of paused Pol II throughout the embryo. Precise gene expression requires both low variability when the gene is active and low background expression when the gene is inactive. Since neither promoter type fulfills both criteria, there may be a tradeoff between them. Highly paused promoters are more optimal for achieving low expression variability when genes are active, while TATA promoters are more optimal for achieving very low background expression when the genes are inactive. This idea is consistent with our finding that these two promoter types exist in a continuum and that promoters that we classified in between (dual or paused promoters) represent different tradeoffs between the two properties.

How paused Pol II changes the expression characteristics of promoters is not entirely clear, but most likely this occurs indirectly, by keeping the promoter open, rather than by promoting activation through formation of the pre-initiation complex (Gilchrist et al. 2010; Shao and Zeitlinger 2017; Gilchrist et al. 2008). We therefore favor a model in which paused Pol II lowers the activation barrier for the promoter by keeping the promoter nucleosome away and increasing the accessibility to the promoter. This model is consistent with our analysis of nucleosome occupancy by MNase-seq and chromatin accessibility by ATAC-seq across our promoter types, showing that paused promoters have significantly higher accessibility than TATA promoters.

A significant role for the promoter nucleosome in shaping the expression characteristic of a gene is supported by studies in yeast. TATA promoters are typically occluded by a promoter nucleosome, which is then removed during gene activation (Tirosh and Barkai 2008; Lee et al. 2007; Tirosh et al. 2007). The stochastic nature of nucleosome removal has been shown to be associated with gene expression variability (Boeger et al. 2008; Kim and O’Shea 2008; Brown et al. 2013; Boeger et al. 2015). Paused promoters in *Drosophila*, on the other hand, also have a promoter nucleosome, but this nucleosome is absent when paused Pol II is present (Gilchrist et al. 2010; Gilchrist et al. 2008; Gaertner et al. 2012), which may lower the activation barrier and be associated with lower gene expression variability.

While the different expression characteristics can explain why effector genes use different promoter types, a not mutually exclusive possibility is that the two promoter types play different roles in evolution. Effector genes in differentiated tissues are under evolutionary pressure to adapt to a changing environment over time. TATA promoters may be more tunable in their expression levels since they are more sensitive to mutational perturbations and show higher expression divergence between species (Hornung et al. 2012; Tirosh et al. 2006; Tirosh and Barkai 2008; Sigalova et al. 2020; Landry et al. 2007). Consistent with this hypothesis, we found that TATA effector genes were often short genes in clusters that are expressed in tissues that have to adjust to their environments, such as the epidermis, gut and trachea. These genes may correspond to the “gene batteries” described by Eric Davidson and their regulation may be relatively simple (Erwin and Davidson 2009). In contrast, promoters with paused Pol II tend to be found in relatively long genes with extensive cis-regulatory regions (Zeitlinger and Stark 2010; Sigalova et al. 2020). Evolutionary adaptations therefore occur more likely in distal cis-regulatory regions since promoter mutations that affect transcription would be likely pleiotropic.

The characteristics of effector genes that we observed for the late *Drosophila* embryo here are likely to be similar in vertebrates. Vertebrate TATA promoters are also enriched among the tissue-specific genes (Schug et al. 2005; Carninci et al. 2006; FANTOM Consortium and the RIKEN PMI and CLST (DGT) et al. 2014) and have higher expression variability (Zoller et al. 2015; Sigalova et al. 2020; Faure et al. 2017). In addition, the presence of paused Pol II at promoters has been found to correlate with reduced gene expression noise (Day et al. 2016). Thus, the possible tradeoff between various expression characteristics that we observe at different promoter types could be a broadly applicable feature of metazoans.

## Materials and Methods

### Fly stocks

*Oregon-R* embryos were used as wild type samples for the whole embryo experiments. For the INTACT experiments, embryos from fly stocks expressing tissue-specific RAN-GAP-mcherry-FLAG-BirA were used. To generate the fly stocks expressing tissue-specific RAN-GAP-mcherry-FLAG-BirA, UAS RAN-GAP-mcherry-FLAG-BirA lines were crossed with tissue-specific Gal4 driver lines in six different tissues as mentioned in Supplementary Excel 2. The Bloomington stock number for the Gal4 drivers crossed with the UAS lines to obtain the above lines are as follows: Neuron – 8760, Glia – 7415, Trachea – 8807, Epidermis – 7021, Muscle – 27390 and Gut – 110394.

### Embryo collection

Adult flies were maintained in population cages for embryo collections and embryos were collected and matured on apple juice plates at 25°C. For example, 14-17h AED embryos were collected on apple juice plates for 3 h at 25°C in cages and then matured at 25°C for another 14 h in the incubator. Embryos were dechorionated for 1 min with 67% bleach then cross-linked for 15 min with 1.8% formaldehyde (final concentration in water phase). Embryos were flash frozen in liquid nitrogen and stored at −80°C. For ATAC-seq and scRNA-seq experiments, the embryos were processed immediately after dechorionation without crosslinking.

### Isolation of tissue-specific nuclei

Nuclei isolation was performed using previously published protocols with modifications (Bonn et al. 2012; Deal and Henikoff 2011). Nuclei were isolated by douncing 0.5 g of embryos in HBS buffer (0.125 M Sucrose, 15 mM Tris (pH 7.5), 15 mM NaCl, 40 mM KCl, 2 mM EDTA, 0.5 mM EGTA, 2% BSA, protease inhibitors) in a 15 ml dounce tissue grinder followed by filtering the nuclei suspension through two layers of miracloth (Calbiochem, #475855). Nuclei were spun at 500g for 10 min at 4°C, and the supernatant was discarded. The nuclear pellet was resuspended in the HBS buffer and dissociated using a syringe (22.5-gauge needle) 10 times. Nuclei were spun again, and the pellet was resuspended in HBS buffer and incubated with Dynabeads^®^ M-280 Streptavidin beads (Invitrogen, # 11205D) for 30 minutes with end-to-end rotation at 4°C. A magnet was used to separate the bead-bound nuclei, and the beads were washed thoroughly with the HBS buffer.

### ChIP-seq experiments

ChIP-seq experiments were performed as described (Chen et al. 2013) with the following differences: ~100 mg OregonR embryos were used per IP. 5 μg chromatin was used for tissue-specific ChIP-seq experiments. Libraries were prepared according to manufacturer’s instructions. ChIP-seq libraries were prepared from 5-15 ng ChIP DNA or 100 ng WCE input DNA.

### ATAC-seq experiments

ATAC-seq was performed using about 500-2000 embryos of stage 14–17 h AED. Nuclei were isolated by douncing embryos in the HBS buffer as described above. Whole embryo ATAC-seq was performed on OregonR embryos without isolating nuclei from specific tissues. The transposition of the nuclei was performed as described in (Buenrostro et al. 2013), using 2.5 μl Tn*5* transposase, followed by PCR amplification (Nextera DNA Sample Preparation Kit: FC-121-1030, Illumina) and library preparation (the Nextera index kit: FC-121-1011, Illumina). Libraries were purified using Agencourt AMPure XP beads (A63881, Beckman Coulter). Paired-end sequencing was performed on the NextSeq 500 instrument (Illumina). Following sequencing, the chromatin accessibility was calculated by computationally filtering for fragments of size 0-100 bp, which represent small fragments from accessible regions.

### MNase-seq experiments

MNase digestion was performed similarly to previously published protocols (Chen et al. 2013). Briefly, chromatin was extracted from 0.1 mg of *wt* OregonR embryos per replicate, then digested with a concentration gradient of MNase (Worthington Biochemical Corporation #LS004798)for 30 min at 37°C. All samples were run on a gel and the digestion concentration to be sequenced was chosen as previously described (Chen et al. 2013). Libraries were prepared using the NEBNext DNA Library Prep kit following the manufacturer's instructions and then paired-end sequenced on an Illumina HiSeq 2500 sequencing system. The nucleosome-sized fragments (100-200 bp) were selected computationally to analyze the nucleosome occupancy.

### mRNA-seq experiments

Total mRNA was extracted from non-cross-linked embryos using the Maxwell Total mRNA purification kit (Promega, #AS1225) according to the manufacturer’s instructions. PolyA-mRNA was isolated using DynaI oligo(dT) beads (Life Technologies, #61002). Libraries were prepared following the instructions of the TruSeq DNA Sample Preparation Kit (Illumina, #FC-121-2001) and sequenced on the HiSeq 2500 and the Nextseq 500 (Illumina).

### scRNA-seq experiments

scRNA-seq experiments were performed on 14-14.5h AED wildtype OregonR embryos. The isolation of single cells was performed similar to the previously published protocol (Karaiskos et al. 2017), with the following modifications. The embryos were dounced in SFX medium with 0.1% PF68 + 0.1% BSA, which was found to improve the cell viability. The total number of dounces was increased to 120 to improve the isolation of cells from late-stage embryos. Isolated cells were filtered and washed and then resuspended in Schneider’s medium to avoid any interference with droplet formations in the subsequent steps. Resuspended cells were immediately processed in the 10× Genomics instrument with optimal loading at a targeted capture rate of about 6000 cells per run to minimize doublets. RNA isolation and cDNA synthesis and amplification were done according to the manufacturer’s instructions. Libraries were then prepared following the instructions of the TruSeq DNA Sample Preparation Kit (Illumina, #FC-121-2001) and sequenced on the HiSeq 2000. scRNA-seq experiments were performed on two biological replicates on separate days from different cages.

### Sequence alignment

All sequencing reads were aligned to the *Drosophila melanogaster* genome (dm6) using Bowtie (v 1.1.2) (Langmead et al. 2009), allowing a maximum of two mismatches and including only uniquely aligned reads. The sequenced reads were trimmed to 50 bp before alignment. Aligned reads were then extended to the estimated insert size or the actual size for the paired-end libraries. For the bulk mRNAseq samples, the gene expression values were calculated by performing pseudo-alignment using the Kallisto package (Bray et al. 2016). For the scRNA-seq samples, alignment and separations of reads from different cells were done using the Cell Ranger pipeline (v 2.1.1) from 10× Genomics.

### Mapping scRNA-seq data to known tissue types

We obtained the gene expression profiles of about 3500 cells in total from two independent biological replicates. The cells from both the replicates were pooled together for the downstream analysis. The Seurat package (Satija et al. 2015) was used for normalization, clustering and visualization of the scRNA-seq data. The cell-gene expression matrix was normalized by the total expression and scaled by a factor of 10000 and log-transformed. Principal component analysis was performed on highly variable genes. The first 20 principal components were used as input for clustering by the Shared Nearest Neighbor method. The Seurat package was also used to identify the marker genes for each of the clusters.

The tissue of origin for the clusters was identified by comparing the scRNA-seq expression patterns with the *in situ* hybridization profiles from the Berkeley *Drosophila* Genome Project (BDGP) similar to the previously published method (Karaiskos et al. 2017). Briefly, the annotated gene expression profiles for the embryonic stage 13-16 were obtained from BDGP, excluding the ubiquitously expressed genes. The scRNA-seq data was then binarized into ON/OFF values, based on whether the expression values are above/below a threshold. The expression value at the 0.9 quantile for each gene was used as the threshold above which it was considered ON. The results did not vary significantly for a wide range of cutoffs. The Matthews Correlation Coefficient was calculated based on this binarized version of our data versus the binarized BDGP data. Each cell was annotated as the tissue with which it had the maximum correlation. Each cluster was then assigned the tissue to which the largest number of cells were annotated. For ambiguous clusters, the occurrence of known tissue markers was analyzed and clusters were manually merged or separated such that they better matched anatomical structures. For example, when more than one cluster was annotated with the same tissue type and we could not find meaningful differences between them, we merged these clusters. Similarly, when small subgroups with distinct tissue types were found within a cluster, the clusters were separated into multiple sub-clusters.

### % of cells with expression and coefficient of variation calculations

When calculating the percentage of cells with detectable transcripts for each gene in each tissue, the tissue with maximum expression was considered as the expressing tissue and the five least expressing tissues were considered as other tissues. The coefficient of variation was calculated as the ratio of the standard deviation of expression divided by the mean expression in the expressing tissue. Only the cells with detectable transcripts were considered for this calculation.

### Promoter element enrichment

In Figure 2C and S3C, the presence of known *Drosophila* promoter elements in each promoter is identified with zero mismatches, in a specified window relative to the TSS (Supplementary Excel 3). For each gene group and each promoter element, the enrichment was calculated as the fraction of genes in a group with a promoter element over the fraction of all genes with the same promoter element. The statistical significance was calculated with Fisher's exact test after correcting for multiple testing by the Benjamini–Hochberg method.

### Pausing index calculations

Pausing index in Fig 2E was calculated as the amount of Pol II ChIP-seq signal in the 200bp window downstream of the TSS divided by the Pol II signal in the 200bp-400bp region downstream from the TSS in the gene body.

### Statistical significance calculations

P values in Figures 2E, 3B, 3C, 3F, 4D, S5A, S5B, S6C, S6D were calculated with the two-sided Wilcoxon test. P values in Figures 2C, S3C, S6B were calculated with the Fisher's exact test with multiple-testing correction, *P < 0.05. P values in Figures S3A and S3B were calculated using the hypergeometric test.

## Data and software availability

Raw and processed data associated with this manuscript have been deposited in GEO under session number GSE120157, which will be available after peer reviewed publication. All data analysis performed in this paper, including raw data, processed data, software tools, and analysis scripts are available through a publicly accessible Amazon virtual machine image. The ami-id will be available after peer reviewed publication. The analysis code is also available on GitHub at https://github.com/zeitlingerlab/Ramalingam_promoter_types_2020.

## Author Contributions and Notes

M.N, V.R, and J.Z conceived the study and designed the experiments. M.N and V.R carried out the INTACT experiments. V.R performed the scRNA-seq experiments. V.R and J.J analyzed the data. V.R, J.Z, M.N, and J.J contributed to data interpretation. V.R and J.Z wrote the manuscript.

The authors declare no conflict of interest.

This article contains supporting information online.

## Acknowledgments

We thank Dr. Steve Henikoff for sharing the w1118; p[UASRG]5/CyO, p[twi-GAL4] p[UAS-EGFP] and w1118; p[UASRG]6 fly stocks, which were used to generate the tissue-specific tagged RAN-GAP expressing fly lines. We thank Kate Hall, Allison Peak and Ana Pinson for technical assistance with the scRNA-seq experiments. We thank Robb Krumlauf, Mounia Lagha, Viraj Doddihal, Zainab Afzal, Charles McAnany, Kaelan Brennan, Khyati Dalal, Curtis Bacon, and Melanie Weilert for their feedback on the manuscript. We thank the Finkelstein lab for the biorxiv template from their github repository.

---

**Figure S1:**
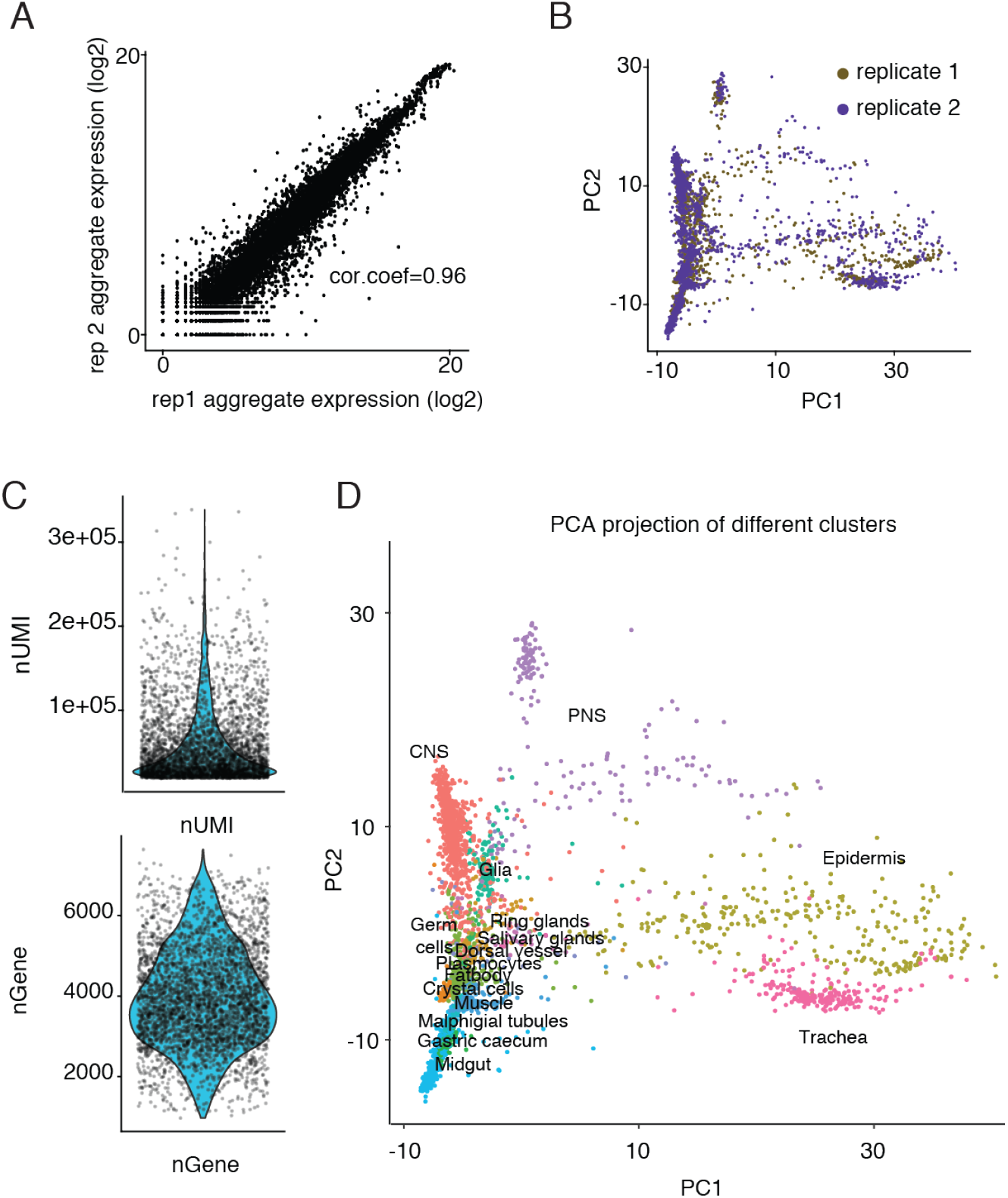
scRNA-seq experiments provide reproducible and high-quality gene expression profiles. (A) A high Pearson correlation coefficient of gene expression levels was observed between single-cell replicates, computed as the sum of read counts for each gene across all cells in both replicates. (B) PCA projection of the scRNA-seq data from both replicates shows the consistency between the two replicates. (C) The number of Unique Molecular identifiers (UMI) and the number of genes captured per cell are shown. On average, about 4000 genes per cell were captured. (D) PCA projection of the scRNA-seq data shows that tissues of similar origin cluster together, which indicates that the main sources of variations in the scRNA-seq data are biologically significant.

**Figure S2:**
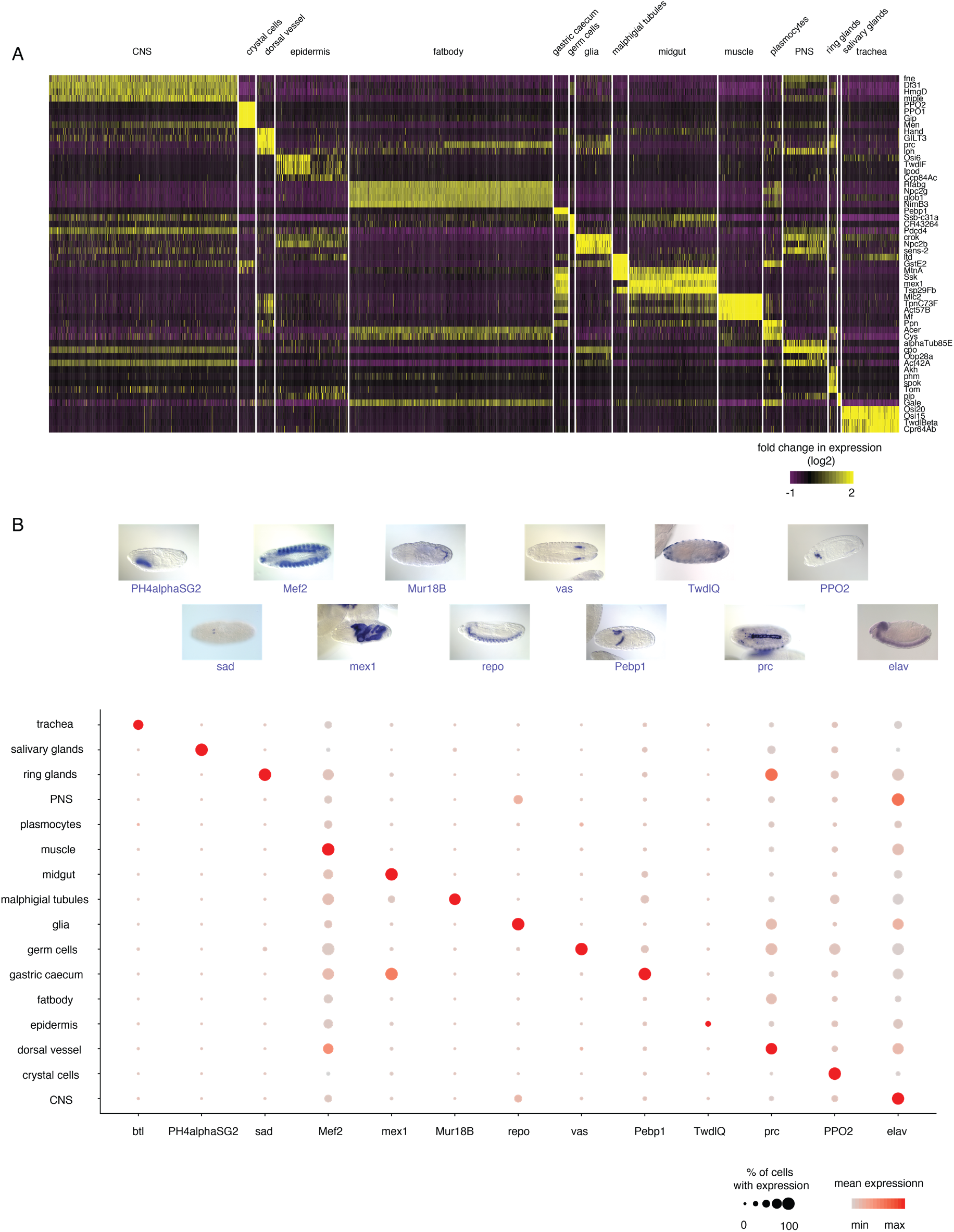
Expression profiles of marker genes of the clusters identified in the scRNA-seq data. (A) Heat map of *de-novo* identified marker genes. The top four differentially expressed genes in each tissue (Wilcoxon test with P value < 0.01), expression in at least 5% of cells in at least one group) is shown. (B) Among them are many known marker genes based on previous studies and *in situ* hybridization from BDGP, as shown in the dot plot. The size of the dot represents the frequency of cells in a tissue with expression and the color intensity represents the mean expression in the tissue.

**Figure S3:**
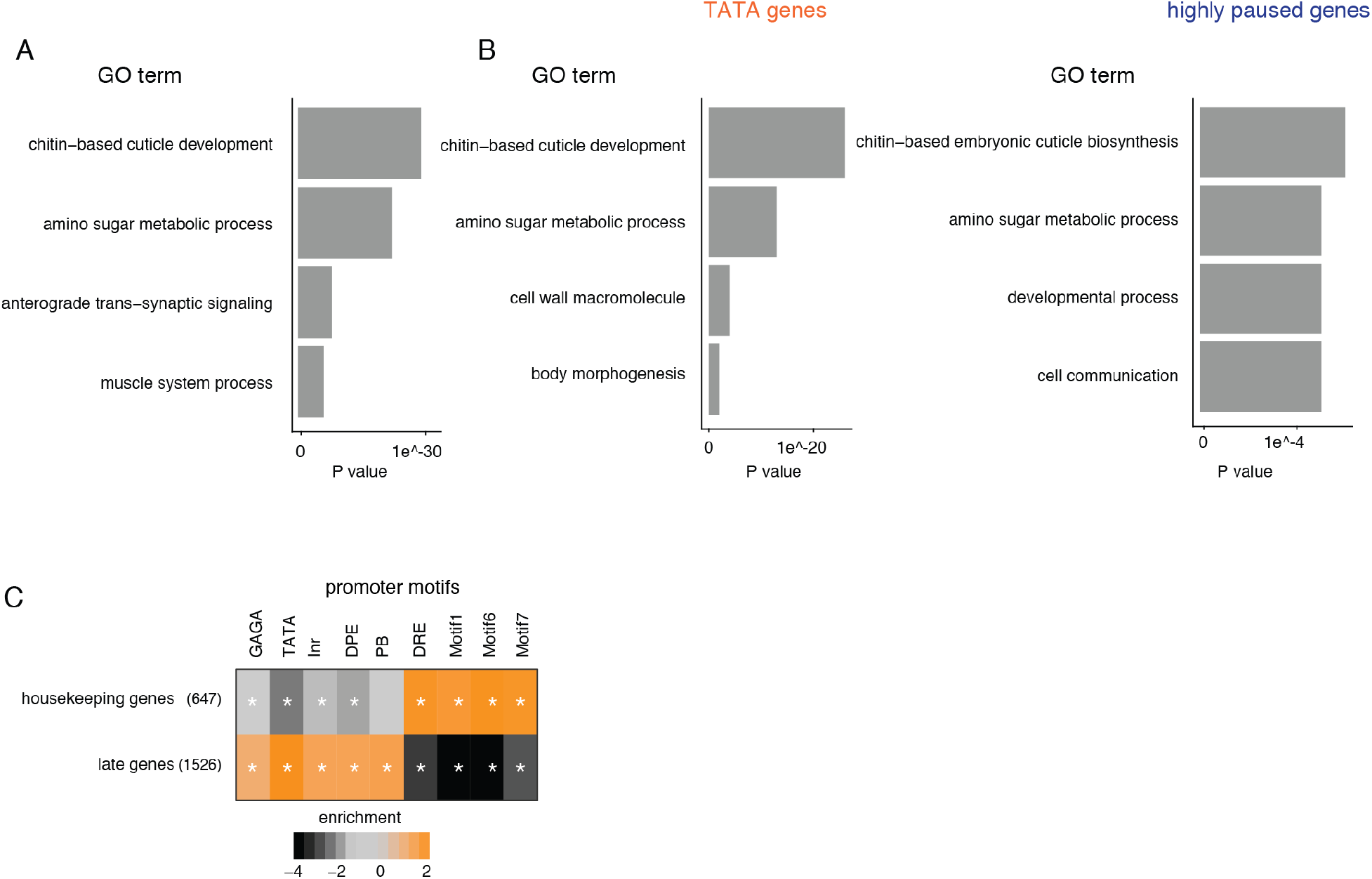
Functional categories and promoter elements of effector genes. (A & B) GO term enrichments were calculated for Biological Processes. For each gene group and each GO term, the enrichment was calculated as the fraction of genes in a group associated with a GO term over the fraction of all genes with the same GO term. The statistical significance for the enriched GO terms was calculated using the hypergeometric test. GO terms associated with less than five genes were not included in the analysis. Top representative terms are shown in the bar plot. (A) Identified effector genes are enriched for expected GO terms. (B) GO term enrichments for the TATA group and the highly paused group effector genes were calculated separately. While there are some differences between the groups, both groups have functional categories expected for effector genes. (C) The identified effector genes were enriched for promoter elements found at tissue-specific genes (focused promoter elements) and are depleted for motifs associated with housekeeping genes (broad promoter elements). A star denotes significance with a Fisher’s exact test, (P <0.05).

**Figure S4:**
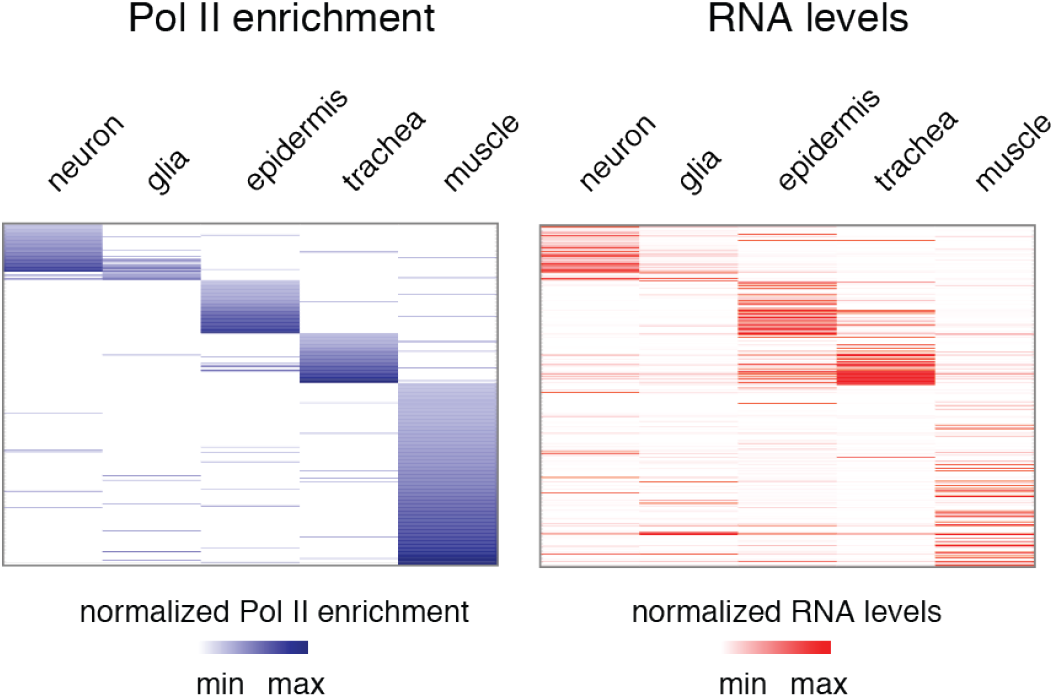
Tissue-specific Pol II ChIP-seq shows specificity and correlates with the scRNA-seq expression profile. The specificity of the tissue-specific Pol II ChIP-seq was evaluated by analyzing the correlation between the Pol II occupancy pattern for a gene and its corresponding expression profile in the scRNA-seq data. The analysis was restricted to genes with Pol II occupancy in a single tissue. The Pol II ChIP-seq signal was calculated around a window starting from the TSS and ending 200bp downstream. There is a clear correspondence between the Pol II occupancy and the scRNA-seq gene expression profile.

**Figure S5:**
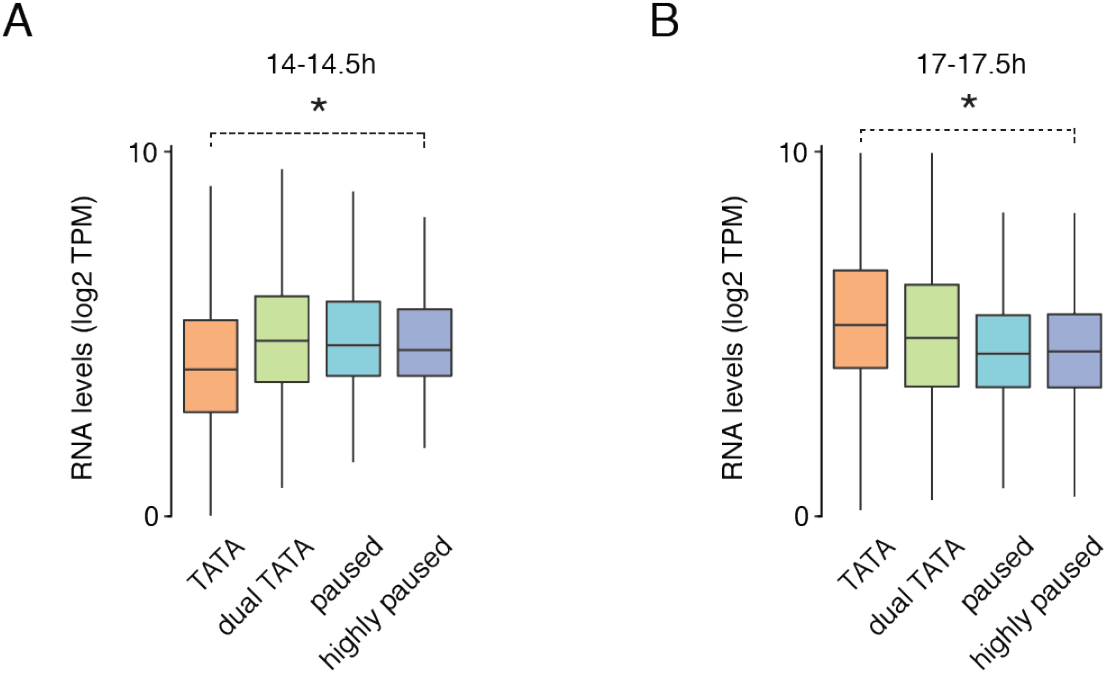
TATA gene expression increases over time. (A & B) RNA levels (log2 TPM) from (A) 14-14.5 h and (B) 17-17.5 h embryos at the different effector gene groups are shown. (A) The TATA genes are expressed at levels lower than the paused genes at 14-14.5 h (Wilcoxon two-sided test, *P <10^−11^). (B) By 17-17.5 h, the TATA genes are expressed at levels higher than the paused genes (Wilcoxon two-sided test, *P <10^−9^).

**Figure S6:**
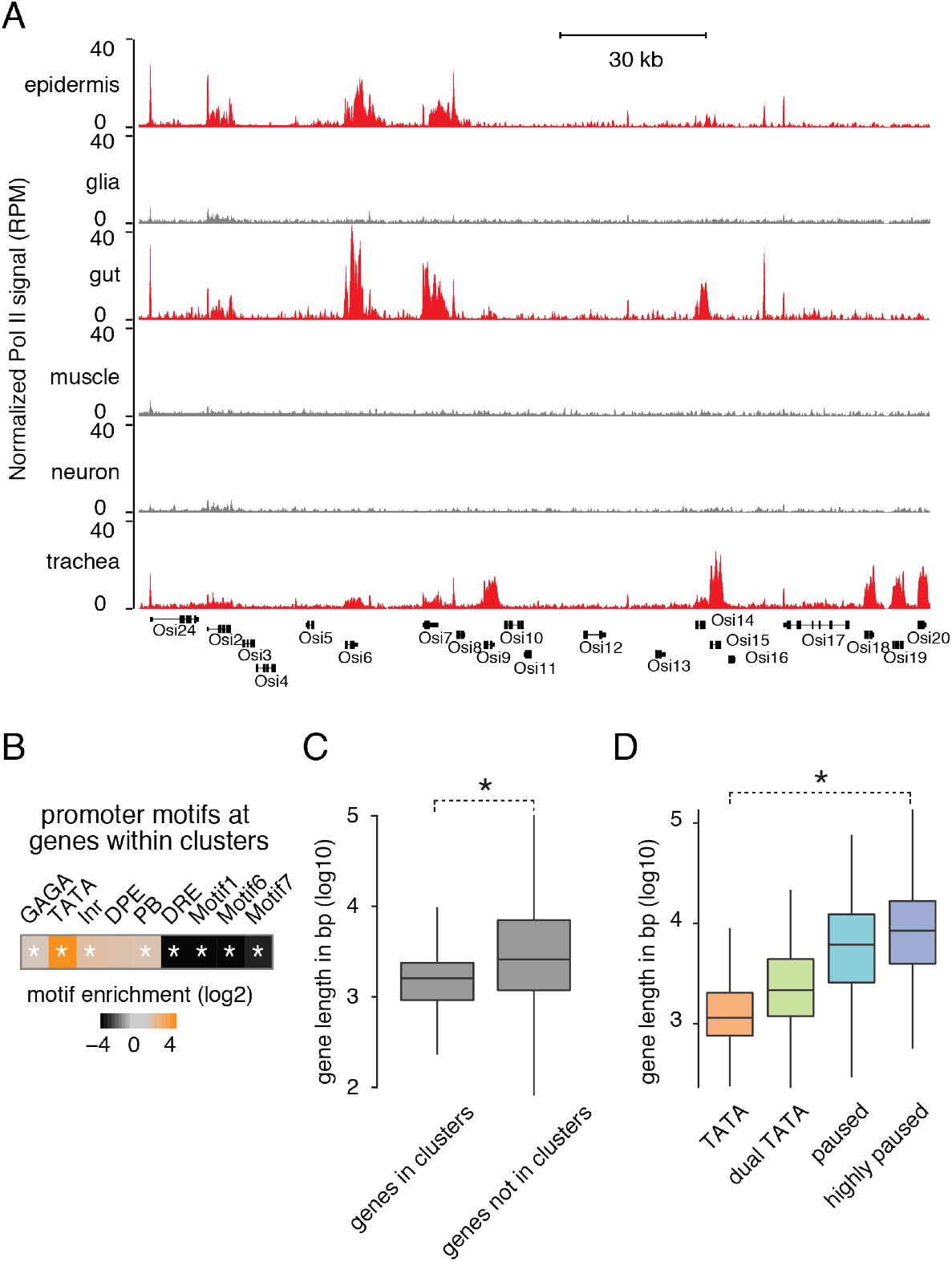
TATA genes often occur in clusters and are short. (A) Read normalized Pol II signals (RPM) at a gene cluster are shown. (B) Promoter elements enriched at genes present in clusters were compared to all other genes. Genes present in clusters are enriched for the TATA motif (*P < 0.05). (C) Total gene lengths of the genes that are present in clusters were compared to all other genes. Genes present in clusters are shorter than the genes which are not present in clusters (Wilcoxon two-sided test, *P <10^−15^). (D) Total gene lengths of genes from different effector gene groups are shown. TATA genes are generally shorter than highly paused genes (Wilcoxon two-sided test, *P <10^−15^).

**Figure S7:**
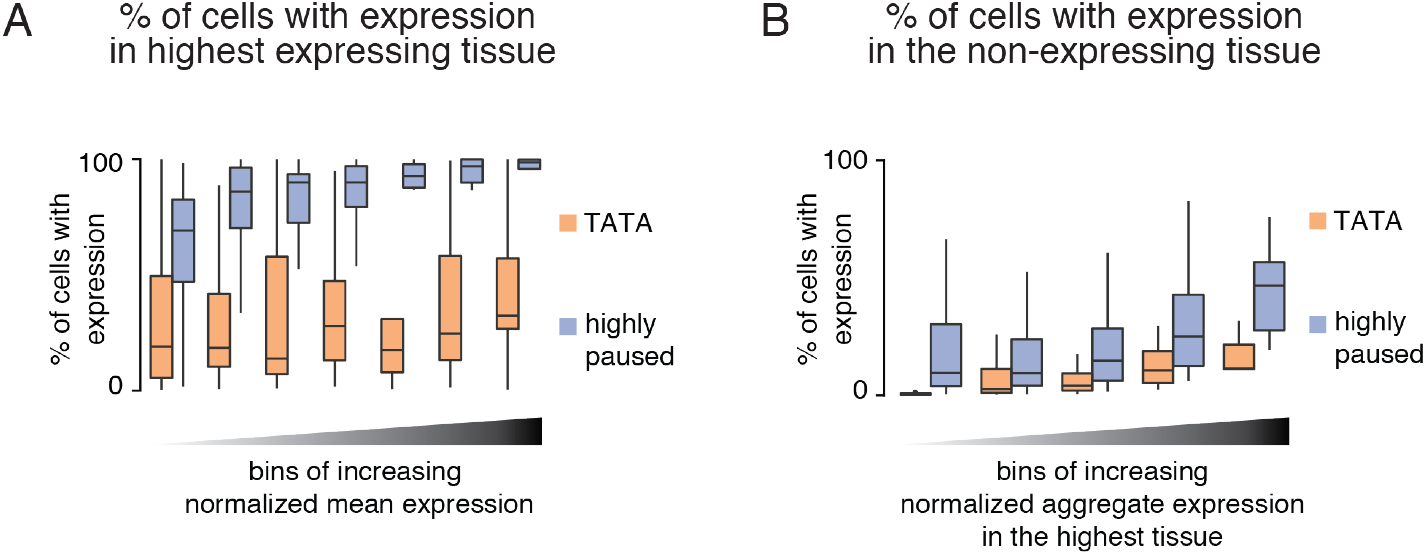
Differences in the scRNA-seq expression profiles of effector gene groups. (A) The coefficient of variation (standard deviation/mean) of gene expression was calculated for putative effector genes in the TATA versus highly paused promoter group, stratified by mean gene expression levels. In all expression bins, the median coefficient of variation was lower for the paused genes compared to the TATA genes. (B) The frequency of cells with detectable expression in the five least expressing tissues (different for each gene) was calculated for all genes in the different effector gene groups, stratified by total expression in the expressing tissues for each gene. The median frequency of expressing cells was higher for the highly paused genes compared to the TATA genes.

